# HLA Allele Imputation with Multitask Deep Convolutional Neural Network

**DOI:** 10.1101/2021.06.03.447012

**Authors:** Calvin Chi

## Abstract

**Motivation:** The Human leukgocyte antigen (HLA) system is a highly polymorphic gene complex encoding the major histocompatibility complex proteins in humans. HLA alleles are of strong epidemiological interest for their large effect sizes in associations with autoimmune diseases, infectious diseases, severe drug reactions, and transplant medicine. Since HLA genotyping can be time-consuming and cost-prohibitive, methods to impute HLA alleles from SNP genotype data have been developed, including HLA Genotype Imputation with Attribute Bagging (HIBAG), HLA*IMP:02, and SNP2HLA. However, limitations of these imputation programs include imputation accuracy, computational runtime, and ability to impute HLA allele haplotypes.

**Results:** We present a deep learning framework for HLA allele imputation using a multitask convolutional neural network (CNN) architecture. In this approach, we use phased SNP genotype data flanking ±250 kb from each HLA locus to simultaneously impute HLA allele haplotyes across loci *HLA-A, -B, -C, -DQA1, -DQB1, -DPA1, -DPB1*, and -*DRB1*. We start by tokenizing phased genotype sequences into k-mers that serve as input to the model. The CNN architecture starts with a shared embedding layer for learning low-dimensional representations of k-mers, shared convolutional layers for detecting genotype motifs, and branches off into separate densely-connected layers for imputing each HLA loci. We present evidence that the CNN used information from known tag SNPs to impute HLA alleles, and demonstrate the architecture is robust against a selection of hyperparameters. On the T1DGC dataset, our model achieved 97.6% imputation accuracy, which was superior to SNP2HLA’s performance and comparable to HIBAG’s performance. However, unlike HIBAG, our method can impute an entire HLA haplotype sequence instead of imputing one locus at a time. Additionally, by separating the training and inference steps, our imputation program provides user flexibility to reduce usage time.

**Availability:** The source code is available at https://github.com/CalvinTChi/HLA_imputation

## 1 Introduction

The major histocompatibility complex (MHC) harbors the human leukocyte antigen (HLA) system on chromosome 6p21.3. HLA genes encode cell-surface proteins that present antigen peptides for recognition by T cells of the host immune system, and are thus among the most polymorphic genes in the human genome [12]. These genes are of strong epidemiological interest due to their large effect sizes in autoimmune diseases, infectious diseases, severe drug reactions, and transplant medicine [4,7,10,25].

Direct typing of HLA alleles include sequence specific oligonucleotide hybridization, capillary sequencing, and next-generation sequencing, but these approaches are labor-intensive, time-consuming, and expensive [9]. Thus, multiple approaches to impute HLA alleles from single nucleotide polymorphism (SNP) genotype data were developed. These methods include HLA Genotype Imputation with Attribute Bagging (HIBAG), HLA*IMP:02, and SNP2HLA [8,14,34]. A previous comparison of HLA imputation programs concluded that HIBAG and SNP2HLA have higher concordance rates than HLA*IMP:02 in European Americans and African Americans [15]. However, HIBAG performs imputation for each locus independently and thus cannot impute HLA allele haplotypes, which limits its applicability for haplotype association studies [16,22]. Lastly, since all these techniques are based on traditional learning algorithms, their performance tend not to scale as well with amount of data in the same way that deep learning does [27]. Given the growing diversity of HLA alleles, deep learning could help increase the number of alleles that could be imputed accurately when more data is available [34]. Recently, Naito *et al*. independently published an effective multitask CNN architecture for HLA imputation that is very similar to the one proposed in this work [26]. This work confirms the effectiveness of this deep learning approach, and provides an additional occlusion analysis.

We present a deep learning approach to HLA imputation using convolutional neural networks (CNN). Deep learning is characterized by its high model capacity to fit arbitrarily complex functions [21]. CNNs have already been successfully applied to a variety of genetic sequence modeling problems, such as DNA-protein binding or chromatin accessibility prediction [24, 33]. CNNs are effective at modeling genetic sequences by learning to detect motifs with its convolutional filters, each of which can be thought of as learning some position weight matrix for a motif [19]. Our multiple input, multiple output CNN accepts phased haplotypes ±250kb flanking each HLA locus and outputs a probability distribution over alleles for each locus. In other words, it maps a haplotype of genotypes to a haplotype of HLA alleles. In this manner, the CNN is able to learn from long range linkage disequilibrium patterns across HLA loci. The embedding and convolutional layers are shared across loci for extraction of higher-order features from flanking genotypes, and then CNN branches off as separate fully-connected layers for each locus. We train and test our model on individuals of European ancestry from the Type 1 Diabetes Genetics Consortium (T1DGC) [29]. We report that our CNN has improved imputation accuracy over SNP2HLA while having comparable performance with HIBAG, but can take considerably less time when the data has already been phased.

## 2 Materials and Methods

### 2.1 T1DGC data

The T1DGC dataset comprises of 5,225 unrelated individuals of European ancestry, with phased genotype and HLA allele data. We start with 5,698 SNPs in the genotype data at the MHC region assayed with the Illumina 550K platform and extract SNPs flanking ±250kb from each HLA locus as predictive SNPs. HLA alleles were typed for *HLA-A, -B, -C, -DQA1, -DQB1, -DPA1, -DPB1* and -*DRB1* at four-digit resolution, totaling 296 distinct alleles. A total of 109 individuals were removed for not having information for both alleles per HLA locus, resulting in 5,116 individuals. Details of this dataset are described elsewhere [14,15,29].

We tokenize, or split, a haplotype sequence of SNPs corresponding to each HLA locus into *k*-mers, with *k* = 5. For example, tokenization of the haplotype sequence AGTCGATAGCAT with *k* = 5 is the process AGTCGATAGCAT → [AGTCG, ATAGC], with the remaining SNPs that cannot form a complete *k*-mer at the right end omitted. Let 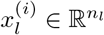 denote the sequence of *n_l_* k-mers corresponding to HLA locus *l* from haplotype *i*. Then haplotype *i* across all eight HLA loci is denoted

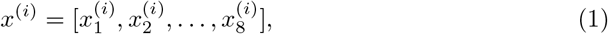

with corresponding HLA alleles 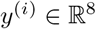 at the four-digit resolution. With m individuals, the dimensions of the dataset are 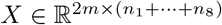 and 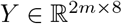, with each individual contributing two haplotypes per HLA locus. The goal of HLA imputation is to find *f* for the mapping *f*: *X* → *Y*.

### 2.2 Data pre-processing

To encode the input haplotypes, one-hot encoding is applied to *n_kmer_* distinct *k*-mers present in the training dataset, with the zero vector serving as a placeholder for unobserved *k*-mers. One-hot encoding involves assigning a 1-to-1 mapping between each *k*-mer to a vector of length *n_kmer_*, with one at the index corresponding to the *k*-mer and zero elsewhere. For instance, when *k* = 1, there are 4 possible *k*-mers (i.e. A, T, C, G), and the one-hot encoding for the third *k*-mer is [0 0 1 0].

Since the embedding and convolutional layers are shared between HLA loci, each haplotype at a HLA locus 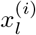 is post-appended with zero vectors to the maximum sequence length *n_max_* = max_*l*_ *n_l_*, such that each HLA locus has the same input feature length of *n*_max_. Thus, the one-hot encoding of a haplotype at HLA locus *l* has dimensions 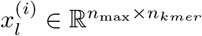, where *n*_max_ is the maximum number of *k*-mers across HLA loci and *n_kmer_* is the *k*-mer encoding length. Each haplotype *i* has multiple inputs 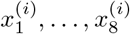 into the CNN, which imputes HLA alleles 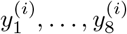.

### 2.3 Network architecture

The CNN architecture is organized into an embedding layer followed by two convolutional layers that is shared between HLA loci. After the convolutional layers, the CNN architecture branches out into separate fully-connected layers for each HLA locus. Overview of the multiple input, multiple output CNN architecture is presented in Figure 1 and details are outlined below

1. Embedding layer of dimension *d* = 8
2. 1D convolution with 64 filters with window size *h* = 4, stride step size *s* = 1, ReLU, 1D max-pool over window of size 4
3. Batch normalization with momentum 0.8, 1D convolution with 64 filters with window size *h* =8, stride step size *s* = 1, ReLU, 1D max-pool over window of size 4
4. Flatten activation feature maps, batch normalization with momentum 0.8, and dropout with probability *p* = 0.5
5. Concatenate activation feature maps between neighboring loci
6. Fully-connected layer with 32 units, ReLU, and dropout with probability *p* = 0.5
7. Softmax output layer

**Figure 1.**
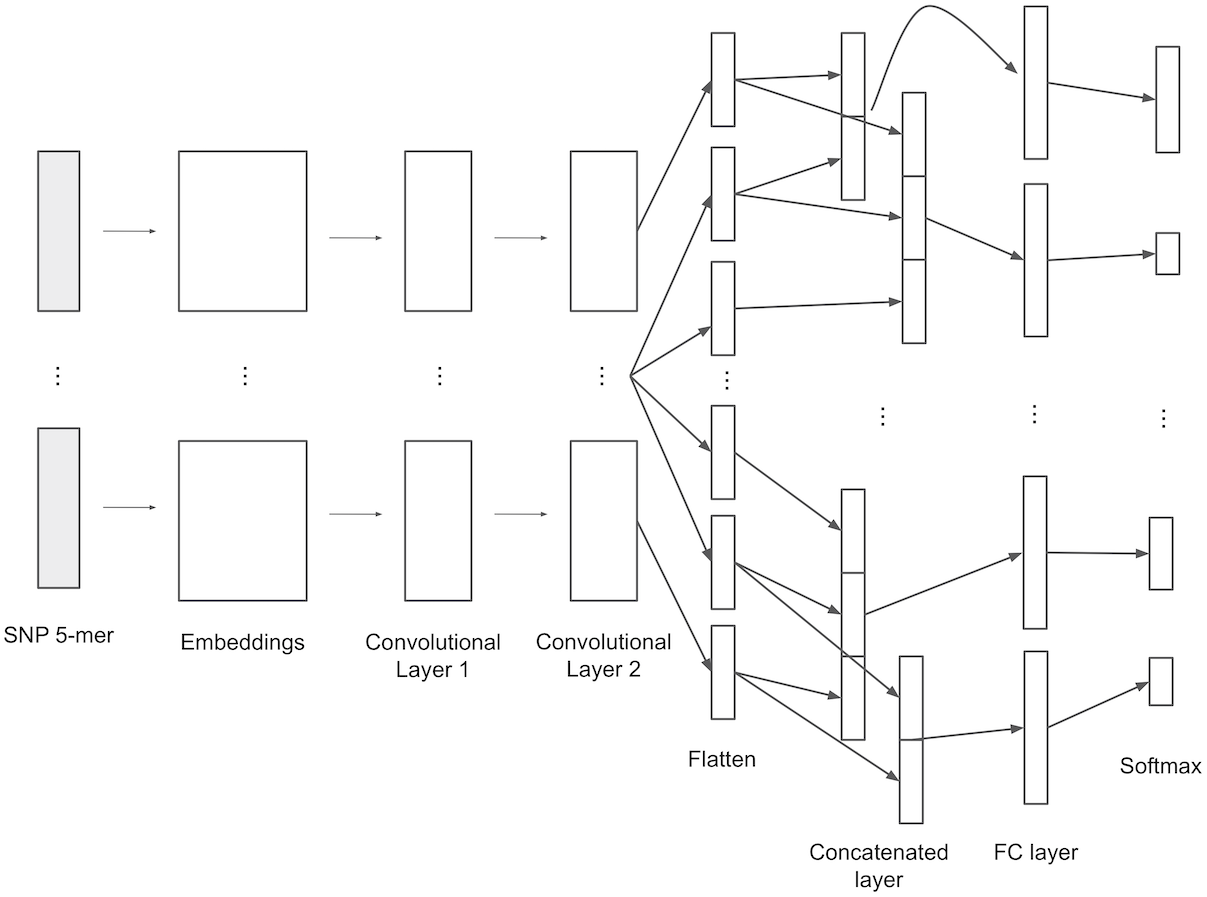
Graphical illustration of CNN architecture for HLA imputation. The *k*-mer sequences are first passed through the shared embedding and convolutional layers to compute intermediate activation feature maps. Activations from neighboring loci are jointly used for imputation by a full-connected network corresponding to each HLA locus. FC = fully-connected.

The first layer of the CNN is an embedding layer for learning a representation of each *k*-mer such that *k*-mers associated with same HLA allele have similar representation. Each *k*-mer is represented by a real-valued vector of dimensions that is typically much less than that for one-hot encoded vectors, which have dimension equal to the number of possible *k*-mers. The purpose of the embedding layer is to transform sparse high-dimensional input vectors into dense low-dimensional vectors that may encode similarities between *k*-mers for improved learning [11]. This concept has been applied in natural language processing tasks [17, 23] as well as prediction task for genetic sequences [24]. The embedding layer is a learnable matrix of dimension (*n_kmer_* + 1) × *d*, where *d* is the size of the embedding dimension and *n_kmer_* + 1 is the number of distinct *k*-mers plus one for the placeholder zero vector. Each row 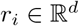 of the embedding matrix can be thought of as the learned representation for the *i*-th *k*-mer.

In our CNN architecture, the embedding layer is followed by two convolutional layers. Each convolutional layer comprises at least the following sequential operations.

1. 1D convolution
2. ReLU nonlinearity
3. Max pooling

The second convolutional layer is preceded by batch normalization. Batch normalization involves normalizing to each input feature independently to have mean zero and variance one, where mean and variance estimates are estimated from mini-batches of data used for stochastic gradient training. To improve representation power of the network, batch normalization also introduces a pair of learnable parameters for scaling and shifting each feature following normalization. The main purpose of batch normalization is to lead to faster training that is less sensitive to parameter initialization. Additionally, batch normalization is also known to have a slight regularization effect [13].

For haplotype i at HLA locus *l*, omit *i, l* for clarity and let 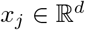 be the *j*-th *k*-mer vector for haplotype *i*. The 1D convolution operation involves applying a trainable filter 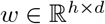 to a window of *h k*-mer vectors to produce a new feature *z_j_* = 〈*w, x_j:j+h−1_*〉_*F*_ + *b*. Here 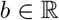 is a learnable bias term and 〈*A, B*〉_*F*_ denotes the Frobenius inner product between matrices *A, B*. The filter *w* is applied against the next window of *h k*-mers in a sliding window fashion with a stride step size *s*. Thus, for haplotype 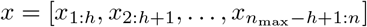, 1D convolution outputs the intermediate feature vector

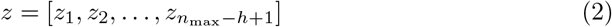

with 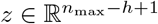. The non-linear ReLU activation function 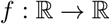 is defined as

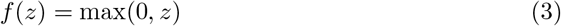

and is applied to each *z_i_* to produce an activation feature *a_i_*. Applying 1D convolution on a window of *h k*-mers followed by ReLU nonlinearity can be succinctly expressed as

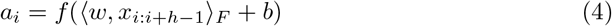

which for *n*_max_ input *k*-mers produces the activation feature vector

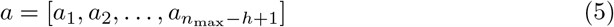

with 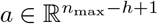. The ReLU activation function is chosen over other non-linearities such as *f*(*x*) = tanh(*x*) or *f*(*x*) = 1/(1 + *e*^−*x*^) because it offers faster training time due to ReLU’s non-saturating property [20]. Finally, max pooling is applied by retaining the maximum activation value over non-overlapping windows of size *p*, reducing the dimension of *a* by *p* and retaining the most important activation feature (i.e. one with largest value in window *h*). Refer to Kim [17] for additional details on the 1D convolution layer.

During training, a filter *w* can be thought of as learning to detect a particular motif across *n*_max_ *k*-mers. A high activation value *a_i_* from filter *w* is indicative of the presence of a motif in window *i*. A convolutional layer learning to detect *f* different motifs would involve training *f* different filters. With *f* filters, the first convolutional layer would have input and output dimensions 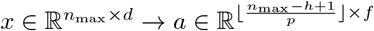. We call this output the activation feature map. In a deep neural network, multiple such convolutional layers can be stacked, with the output activation feature map from the previous layer serving as input to the next layer.

The final portion of our CNN architecture is a fully-connected (FC) layer for final prediction. Following convolution, activation feature maps corresponding each HLA loci *a*_1_,… *a*_8_ are first flattened, then neighboring activations are concatenated according to

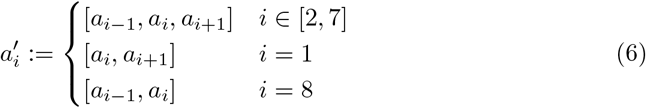

so that our CNN can learn from long-range disequilibrium patterns between neighboring loci. For example, since *HLA-DRB1*15:01* and *HLA-DQB1*06:02* are strongly linked in European populations, presence of *HLA-DRB1*15:01* is indicative of the presence of *HLA-DQB1*06:02* [1]. The output layer for each HLA locus is a softmax layer that outputs a probability distribution over alleles. For example, for *L* possible alleles at a HLA locus, the softmax function for probability of allele *j* given input feature activations 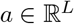 is

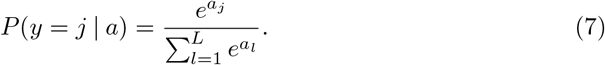

Let 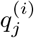 be the predicted probability of the true allele at locus *j* of the *i*-th haplotype, then the target loss function we optimize over is the categorical cross entropy

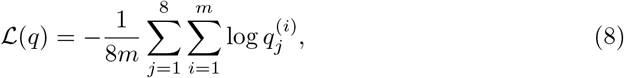

where *m* is number of haplotypes and *q* = {*q*^(1)^,…, *q*^(*m*)^} with 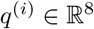.

### 2.4 Regularization

Dropout is applied to the respective activation feature maps feeding into the FC and softmax layers to prevent overfitting [30]. Dropout involves retaining activation values with some fixed probability *p* and zero otherwise, independent of other activation values, with each forward pass during training. With vector inputs, dropout is implemented with a binary mask vector 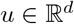 that is multiplied element-wise with the input

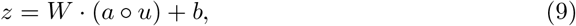

where ∘ denotes element-wise multiplication, 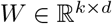 is the trainable weight matrix of a layer, and 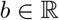 is a trainable bias term. Applying dropout during training can be interpreted as updating weight parameters of a sampled neural networks within the full neural network. By approximating the process of combining exponentially many different neural network architectures, dropout prevents overfitting [30]. At test time, each activation value is multiplied by dropout probability *a*:= *pa* so that the activation value has the same expected output in test and training time.

### 2.5 Hyperparameter optimization and training

We partition 70% of individuals for training and model development, and the remaining individuals for final evaluation. We perform random search over 100 random samples from the hyperparameters embedding dimension, batch size, number of convolutional filters, filter stride step size, number of hidden units for the FC layer, max-pooling window, and dropout probability. Random search is more computationally efficient than exhaustive grid search and can outperform grid search when only a small number of hyperparameters affect final model performance [3]. The set of hyperparameters is optimized over 20% of the training dataset set aside as the validation dataset.

We choose the Adam optimizer for stochastic optimization [18] with a learning rate 0.001 and batch size of 512 haplotypes. During training we apply early stopping when the loss over the development dataset (10% – 15% of the training dataset) does not improve for two epochs, or two passes over the entire training dataset.

## 3 Results and discussion

### 3.1 HLA imputation performance

HLA imputation accuracy on the test dataset was comparable to the state of the art performance achieved with SNP2HLA and HIBAG, which are the HLA imputation programs publicly available to date. Figure 2 and Table 1 summarize the imputation accuracies by imputation program and HLA locus. The overall accuracies for CNN, HIBAG, and SNP2HLA were 97.6%, 97.5%, and 95.8% respectively. Thus, CNN and HIBAG had the best and most comparable performance. Either CNN or HIBAG had superior accuracy over SNP2HLA for every HLA loci. HIBAG out-performed CNN for *HLA-A, HLA-B, HLA-DQA1*, and *HLA-DQB1*. CNN out-performed HIBAG for *HLA-C, HLA-DPA1, HLA-DPB1*, and *HLA-DRB1*.

**Figure 2.**
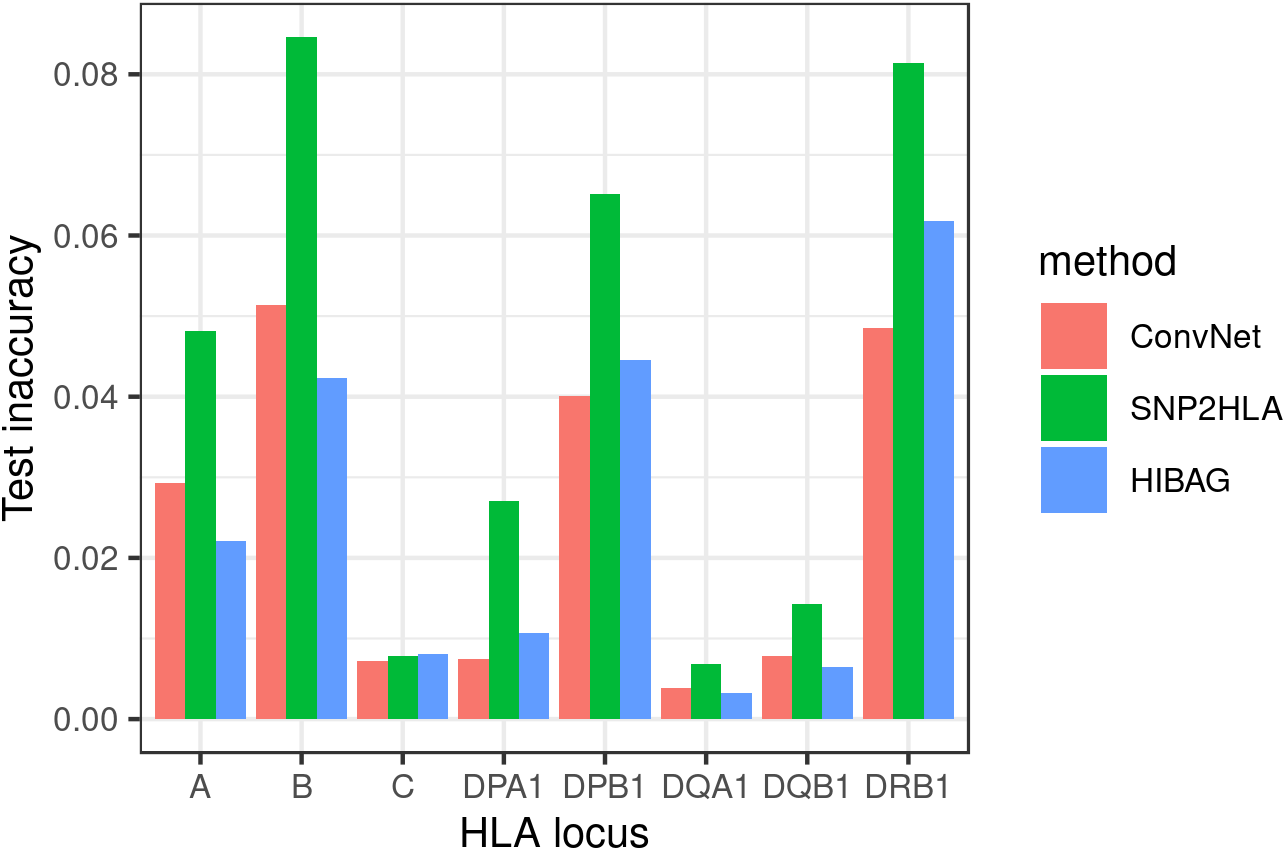
Comparison of test imputation performance by HLA locus between HLA imputation methods. Performance is reported with inaccuracy.

**Table 1.** Comparison of test imputation accuracy by HLA locus between HLA imputation methods.

|  | CNN | SNP2HLA | HIBAG |
| --- | --- | --- | --- |
| *HLA-A* | 0.971 | 0.952 | **0.978** |
| *HLA-B* | 0.949 | 0.915 | **0.958** |
| *HLA-C* | **0.993** | 0.992 | 0.992 |
| *HLA-DPA1* | **0.993** | 0.973 | 0.989 |
| *HLA-DPB1* | **0.960** | 0.935 | 0.955 |
| *HLA-DQA1* | 0.996 | 0.993 | **0.997** |
| *HLA-DQB1* | 0.992 | 0.986 | **0.993** |
| *HLA-DRB1* | **0.951** | 0.919 | 0.938 |
| Overall | **0.976** | 0.958 | 0.975 |

HIBAG out-performed CNN the most at locus *HLA-B* (by about 1%), which is the most polymorphic gene in our training dataset with 96 alleles. It is possible that HIBAG is effective for polymorphic genes due to its design as an ensemble classifier that employs both bootstrap aggregation on individuals and feature bagging on SNPs. However, this does not necessarily imply that deep learning is less effective for imputation of high polymorphic genes, since the size of the T1DGC dataset is relatively small compared that in other deep learning applications.

Our CNN presents an advantage in training time over that required by SNP2HLA and HIBAG when the genotypes are already phased. For HIBAG, 10 classifiers are trained in parallel over 7 CPU cores for each HLA locus. All training was performed by a Linux server with four 10-core Intel(R) Xeon(R) CPU E7-4860 processors (running at 2.8 GHz, with 640KiB/2560KiB/24MiB L1/L2/L3 cache, and using a 64-bit architecture) and a total of 128 GB RAM. The runtimes are presented in Table 2. Both SNP2HLA and HIBAG perform phasing regardless of whether the input is phased or not, which account for the longer times required for imputation. The runtime for HIBAG to impute all eight HLA loci is the longest because imputation is performed separately and independently for each locus. As a consequence, a limitation of HIBAG is its inability to impute HLA haplotypes. It should be noted however, that the runtime of HIBAG could be reduced if imputation for each HLA locus could be performed in parallel. Compared to CNN and HIBAG, which separates the training and testing procedures, the program SNP2HLA requires repeating the entire imputation procedure with the reference panel for each test imputation, which could amount to additional runtime in practice.

**Table 2.** Comparison of imputation program runtimes. HIBAG was trained with 10 classifiers over 7 CPU cores per HLA locus and runtime is summed over HLA loci. SNP2HLA runtime includes test imputation since it combines training and imputation into one procedure. *Since the runtimes of SNP2HLA and HIBAG include phasing, we also report the runtime for CNN that includes phasing performed by BEAGLE in brackets [5].

| Imputation program | Training time (hours) |
| --- | --- |
| CNN | 0.25(9.6)* |
| SNP2HLA | 10.5 |
| HIBAG | 32.3 |

We computed approximate error bars for CNN test accuracy per locus to get an estimate of the variation in imputation accuracy due to variation in test allele frequencies. This is accomplished by creating B = 1, 000 bootstrap samples (with replacement) from the original test dataset and computing the test accuracy from the trained CNN for each bootstrap sample. The distribution of test accuracies are shown in Figure 3.

**Figure 3.**
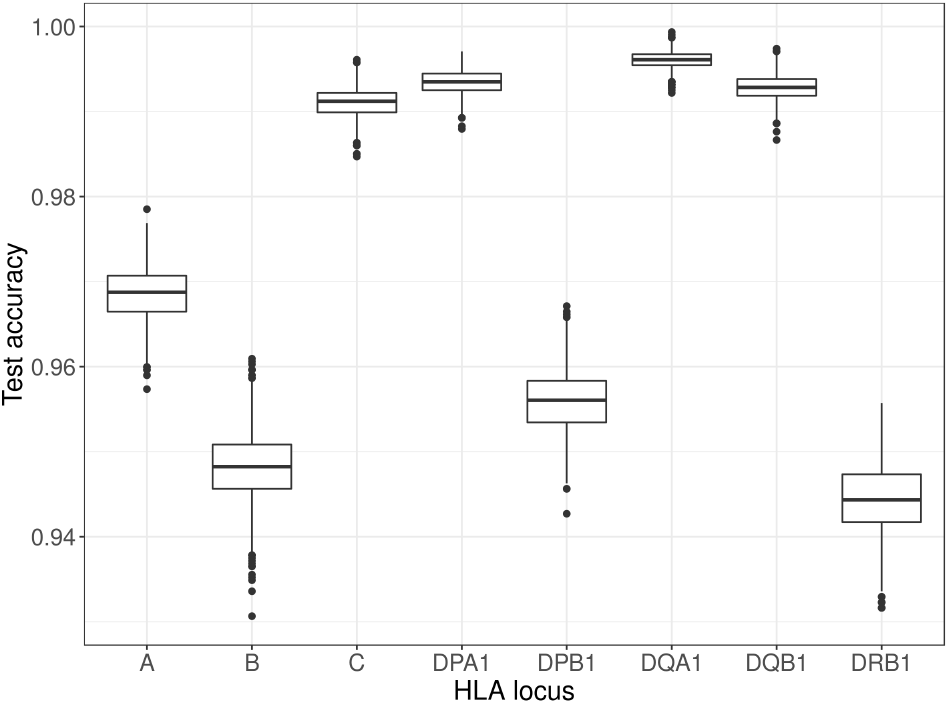
Boxplot of bootstrap test accuracies by HLA locus. The upper whisker extends from the upper hinge (3rd quartile) to the largest value no further than 1.5 the inter-quartile range (IQR) from the upper hinge. The lower whisker extends from the lower hinge (1st quartile) to the smallest value at most 1.5 the IQR of the lower hinge.

The variability in per locus accuracy is correlated with the degree of corresponding polymorphism. The number of HLA alleles in the training dataset is 96 for *HLA-B*, 51 for *HLA-DRB1*, 50 for *HLA-A*, 33 for *HLA-DPB1*, 31 for *HLA-C*, 18 for *HLA-DQB1*, 8 for *HLA-DQA1*, and 7 for *HLA-DPA1*.

### 3.2 Occlusion Analysis

One way to assess what SNPs the CNN might be using to learn the mapping between phased genotypes to HLA alleles is to perform occlusion analysis, which involves independently “masking”, or occluding, a section of SNPs and observing how the probability of the true HLA allele from the trained model subsequently decreases. The larger the decrease in probability, the more important the “masked” SNPs are for imputation. Specifically for a given HLA locus, we successively set 30 *k*-mers corresponding to the locus to zero at a time, which removes information from the selected 30 *k*-mers.

Given we know the tag SNPs for many HLA alleles [6], occlusion analysis can help us determine to what extent the CNN is using the tag SNPs to impute the correct allele. For this analysis we selected the HLA alleles *HLA-DRB1*15:01, HLA-DRB1*04:01, HLA-DQB1*03:02, HLA-DQA1*01:02, HLA-B*08:01*, and *HLA-C*16:01*, many for their association with autoimmune diseases [2]. The allele *HLA-DRB1*15:01* is associated with multiple sclerosis, *HLA-DRB1*04:01* is associated with rheumatoid arthritis, *HLA-DQB1*03:02* is associated with celiac disease, *HLA-DQA1*01:02* is associated with systemic lupus erythematosus, and *HLA-B*08:01* is associated with plasma beta-2 microglobulin [2,31]. Figure 4 shows the resulting probability heatmap from occlusion analysis. There is evidence that the CNN learned to impute the alleles *HLA-B*08:01, HLA-DQB1*03:02, HLA-DQA1*01:02*, and *HLA-DRB1*15:01* based more so on tag SNPs as opposed to other SNPs, since the drops in probability were largest around the respective tag SNPs. This is especially true for *HLA-DRB1*15:01*, where the probability dropped by nearly half when SNP *k*-mers in the neighborhood of tag SNP rs3135388 were “masked”.

**Figure 4.**
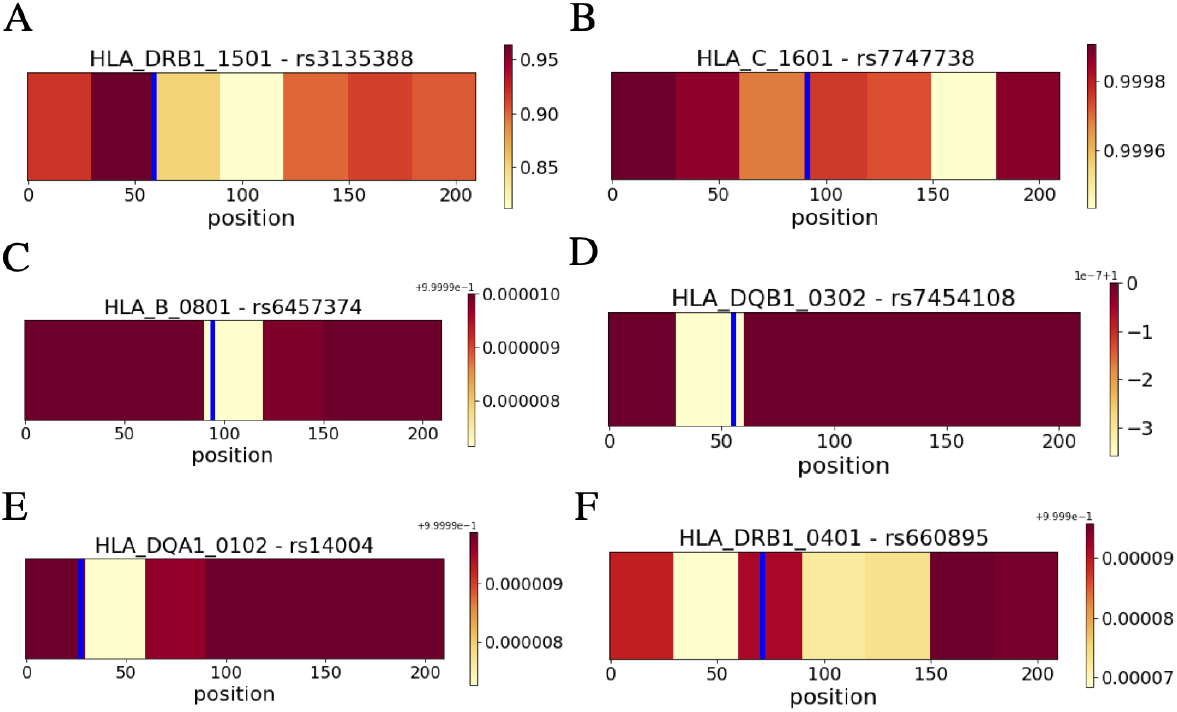
Occlusion sensitivity analysis for HLA alleles (A) *HLA-DRB1*15:01*, (B) *HLA-DRB1*04:01*, (C) *HLA-DQB1*03:02*, (D) *HLA-DQA1*01:02*, (E) *HLA-B*08:01*, and (F) *HLA-C*16:01*. Each panel is a heatmap of true allele probability after blocks of 30 *k*-mers were “masked” to zero for a HLA allele, with the rs number of tag SNPs in the title. The position of the tag SNP is marked with a blue vertical line.

Except for *HLA-DRB1*15:01*, the drops in probability were mostly minuscule, which shows that the CNN learned the imputation mapping based also on genetic information other than what is provided by the tag SNP neighborhood. This suggests that imputation performance by the CNN is robust against genotying errors. For alleles *HLA-C*16:01* and *HLA-DRB1*04:01*, “masking” the neighborhood of tag SNPs did not result in the largest decrease in probability. Together, this demonstrates that the CNN learned to impute based on known associations between SNPs and HLA alleles, but that this was not necessarily the case for all HLA alleles.

### 3.3 Sensitivity analysis

We perform sensitivity analysis to check the robustness of our CNN architecture against some hyperparameters, since the hyperparameter tuning process itself is noisy due to random weight initialization, stochastic optimization, etc. We chose to focus on the hyperparameters *k*-mer length *k*, embedding dimension *d*, and stride step size *s* for all convolutional filters *f*.

According to Figure 5, performance of the CNN is not sensitive to the choices of embedding dimension *d* and *k*-mer length *k*. We tested the embedding dimensions *d* = 5, 10, 15, 20, which controls the model complexity due to the embedding layer. We varied the *k*-mer length for *k* = 3, 4, 5,6, 7, which controls the input dimension to the CNN.

**Figure 5.**
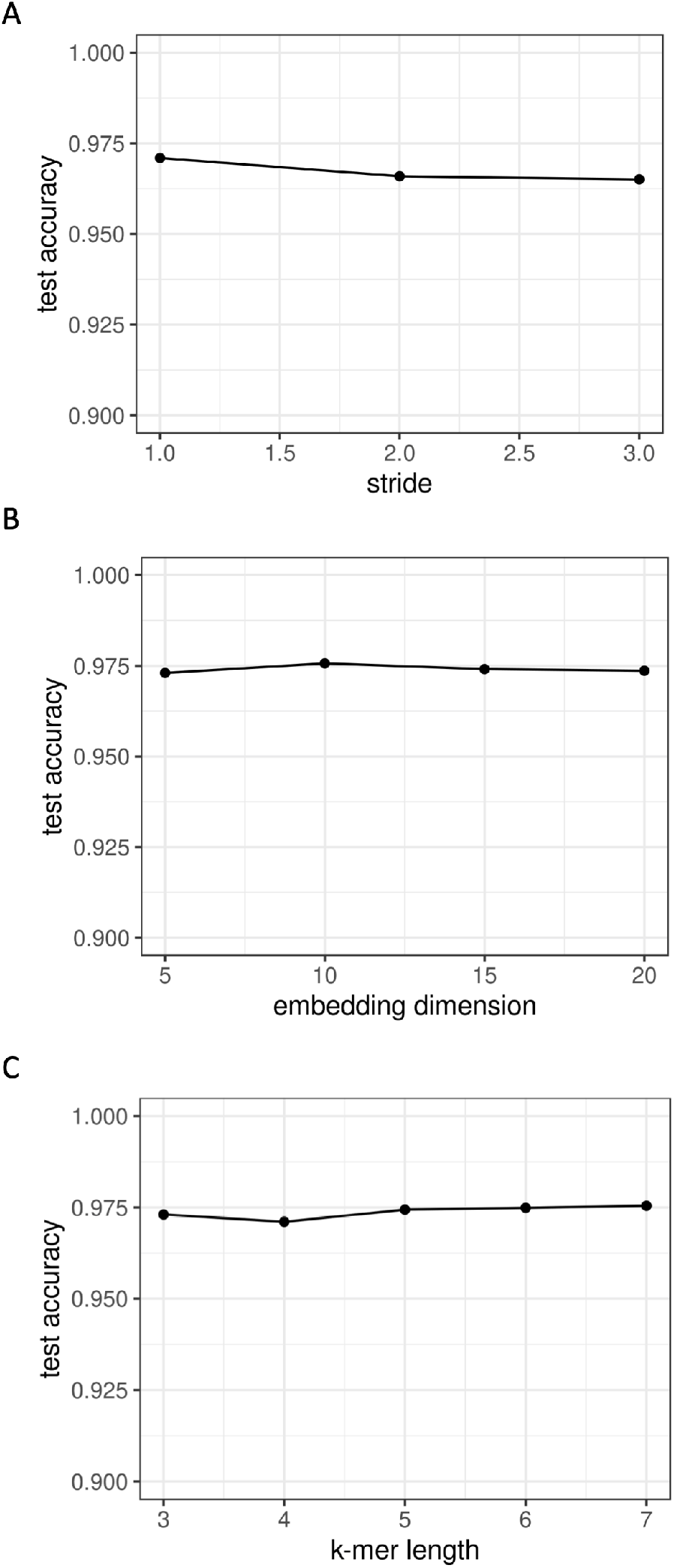
Sensitivity analysis of hyperparameters (A) convolutional filter *f* stride step size *s*, (B) embedding dimension *d*, and (C) *k*-mer length *k*, performed on the test dataset.

However, performance decreased slightly with stride step size *s*, where step sizes *s* = 1,2,3 were tested. This could be due to the fact that a larger step size *s* reduces the dimension of the activation feature maps more aggressively between layers of the CNN and discards more information. Thus, the recommended step size for the CNN is *s* = 1.

## 4 Conclusion

We introduce a 1D CNN that simultaneously imputes alleles at the four-digit resolution for HLA loci *HLA-A, -B, -C, -DQA1, -DQB1, -DPA1, -DPB1* and -*DRB1* from phased genotype data flanking each locus. Genotype data corresponding to each HLA locus are first tokenized into units of *k*-mers and one-hot encoded before serving as input to the CNN. The CNN architecture starts with an embedding layer that learns dense low-dimensional vectors to represent *k*-mers from high dimensional one-hot encodings. Following the embedding layer are two convolutional layers that learn to detect genotype motifs for HLA imputation. The activation feature maps learned for each locus are concatenated with feature maps of neighboring loci for final allele imputation using fully-connected networks. The concatenation of neighboring activations allows genotype information corresponding to a HLA locus to influence imputation of neighboring loci when there exists long-range disequilibrium between neighboring loci.

Our CNN shares the ability of SNP2HLA to impute HLA allele haplotypes while having high imputation accuracy comparable to that from HIBAG. Additionally, separation of the stages of phasing, training, and imputation reduces unnecessary runtime that results from combining these stages into one. We demonstrate via occlusion sensitivity analysis that CNN can indeed learn to impute HLA alleles based on known associations between tag SNPs and HLA alleles, but also that the learned mapping between genotype and HLA allele is dependent on more than these known associations. The architecture we propose is relatively robust against the hyperparameters embedding dimension *d*, stride step size *s*, and *k*-mer length *k*. We anticipate that the performance of deep learning for HLA imputation will improve as more SNP and HLA datasets become available.

Although we demonstrate the effectiveness of CNN for imputation of HLA alleles at the four digit resolution, our CNN can in principle also impute amino acid polymorphisms in HLA proteins from SNP genotype data. A extension of our work is to explore how effective deep learning could be for HLA imputation in admixed populations, which are populations with more than one ancestral populations. A study found that HIBAG was able to achieve reasonable HLA imputation accuracy in admixed Brazilian population using the 1000 Genomes HLA and SNP dataset [28]. However, a decrease in HLA imputation accuracy is generally observed in admixed populations compared to that in European populations [15]. Part of this is due to a partial mismatch between the ancestries present in the reference population and the population to be imputed, but another could be due to insufficient conditioning on population substructure and differing linkage disequilibrium patterns at the MHC between populations. It is known that many HLA alleles are population-specific [32]. One potential approach to take genetic ancestry into consideration is via multi-task deep learning that learns to map SNP genotype data to both HLA alleles and genetic ancestry. A successful application of deep learning for HLA imputation in admixed populations would likely require careful consideration of network architecture as well as how to take into account genetic ancestry.

## Acknowledgements

We would like to thank Professor Haiyan Huang for her suggestions and Professor Lisa Barcellos for her suggestions and computing resources.

